# The temporal structure of the inner retina at a single glance

**DOI:** 10.1101/743047

**Authors:** Zhijian Zhao, David Klindt, André Maia Chagas, Klaudia P. Szatko, Luke Rogerson, Dario A. Protti, Christian Behrens, Deniz Dalkara, Timm Schubert, Matthias Bethge, Katrin Franke, Philipp Berens, Alexander Ecker, Thomas Euler

## Abstract

The retina decomposes visual stimuli into parallel channels that encode different features of the visual environment. Central to this computation is the synaptic processing in a dense and thick layer of neuropil, the so-called inner plexiform layer (IPL). Here, different types of bipolar cells stratifying at distinct depths relay the excitatory feedforward drive from photoreceptors to amacrine and ganglion cells. Current experimental techniques for studying processing in the IPL do not allow imaging the entire IPL simultaneously in the intact tissue. Here, we extend a two-photon microscope with an electrically tunable lens allowing us to obtain optical vertical slices of the IPL, which provide a complete picture of the response diversity of bipolar cells at a “single glance”. The nature of these axial recordings additionally allowed us to isolate and investigate batch effects, i.e. inter-experimental variations resulting in systematic differences in response speed. As a proof of principle, we developed a simple model that disentangles biological from experimental causes of variability, and allowed us to recover the characteristic gradient of response speeds across the IPL with higher precision than before. Our new framework will make it possible to study the computations performed in the central synaptic layer of the retina more efficiently.

## Introduction

The primary excitatory pathway of the mouse retina consists of photoreceptors, bipolar cells (BCs) and retinal ganglion cells (RGCs) (reviewed in refs^1,2^). At the core of this pathway is the inner plexiform layer (IPL), a dense synaptic plexus composed of the axon terminals of BCs, the neurites of amacrine cells, as well as the dendrites of RGCs. Specifically, the photoreceptor signal is relayed by the BCs to the RGCs via glutamatergic synapses (reviewed in ref^3^). This “vertical” transmission is shaped by mostly inhibitory interactions with amacrine cells, which integrate signals laterally along and/or vertically across the IPL (reviewed in ref^4^). Amacrine cells modulate, for instance, the sensitivity of BCs to certain spatio-temporal features^5–7^.

Within the IPL, the axon terminals of each of the 14 BC types^8–12^ project to a distinct depth with axonal profiles of different BC types partially overlapping and jointly covering the whole depth of the IPL^10,11,13^. Functionally, each BC type constitutes a particular feature channel, with certain temporal dynamics^7^, including On and Off BC types sensitive to light increments or decrements, respectively^14^, different kinetics^15,16^, and chromatic signals^17,18^. Some of these features are systematically mapped across the IPL: For example, On BCs project to the inner and Off BCs to the outer portion of the IPL^14,19^. Also kinetic response properties appear to be mapped, with the axonal profiles of more transient BCs localised in the IPL centre^7,15,20,21^.

To study BC function, early studies mostly used single-cell electrical recordings in vertical slices, where many lateral connections (e.g. large-scale amacrine cells) are severed, only electrical signals in the cell body of BCs can be recorded and experiments are time consuming. Recently, two-photon (2P) Ca^2+^ or glutamate imaging in the explanted whole-mount retina has been introduced as a high-throughput alterative^7,15,20^. This approach preserves the integrity of the retinal network, but typically requires recording horizontal planes (x-y scans) at different IPL levels. Since the activity at different IPL depths is recorded sequentially, it can be more difficult to disentangle functional differences between the signals represented at different depths and experimental factors inducing differences between scans.

Here, we introduce fast axial x-z scanning, a method to image across the entire IPL depth near simultaneously in the intact whole-mount retina through “vertical optical slices”, by equipping a 2P microscope (Fig. 1)^22–24^ with an electrically tunable lens (ETL) to quickly shift the focus along the z axis^25^. We evaluated these x-z scans by imaging light stimulus-evoked glutamate release using iGluSnFR^26^ ubiquitously expressed via AAV transduction. We found that axial x-z scans can capture functional signals at different IPL depths similarly well compared to “traditional” x-y scans^7^. Because the new scanning mode allowed us to acquire signals with different response polarity and kinetics within a single scan, we were able to identify and correct for batch effects, corresponding to inter-experimental variation caused by experimental and biological factors like indicator concentration and temperature. As a proof of principle, we show how correcting for batch effects can markedly improve recovering the characteristic change in response kinetics across the IPL, where fast signals are represented towards the middle and slower signals towards the borders of the IPL^7,15,20,27^.

**Figure 1.**
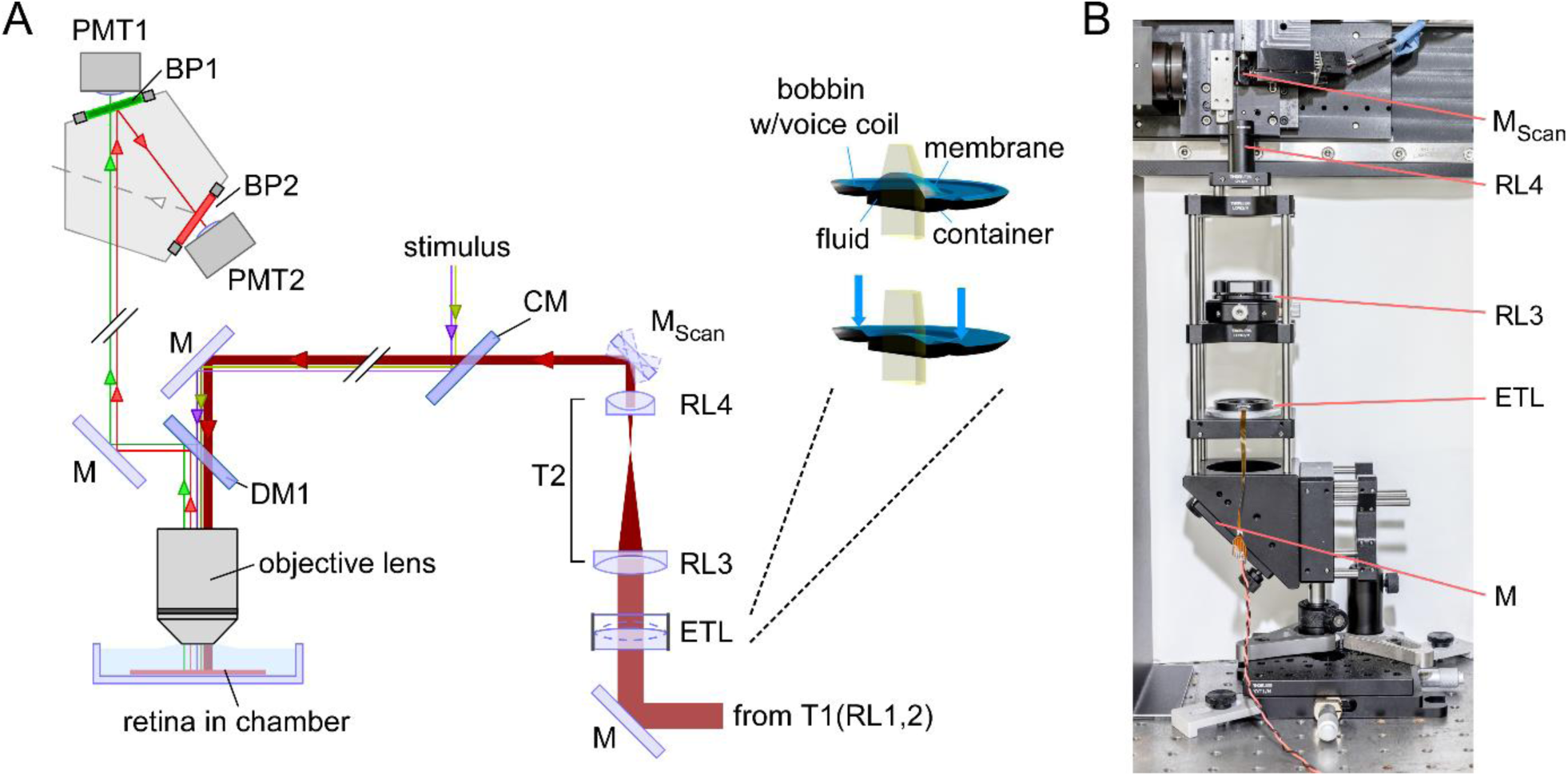
Overview of the two-photon (2P) microscope equipped with an electrically tunable lens (ETL). For simplicity, most lenses and silver mirrors (M) were omitted. For parts, see Table 1. A, Schematic diagram of the microscope’s main optical paths, with the EL-16-40-TC (Optotune) inserted before the x-y galvo scan mirrors (M_Scan_). Inset: Cross section and working principle of the ETL; a voice-coil actuator generates pressure on a container, which in turn pushes optic fluid into the lens volume sealed by polymer membrane and, thereby, modulating the curvature of the lens surface. B, Photograph of the excitation path before the scan mirrors. CM, cold mirror; DM, dichroic mirror; BP, band pass filter; PMT, GaAsP photomultiplier tube; RL, relay lens; T, telescope. Panel A adapted from Euler et al.^22^, inset adapted from Optotune website (https://www.optotune.com/).

## Methods

### Equipping the 2P microscope with an ETL

To allow for axial scanning, we modified a movable objective microscope (MOM, designed by W. Denk, now MPI Martinsried; purchased from Sutter Instruments/Science Products, Novato, CA). Design and configuration of the MOM have been described elsewhere^7,22,23,28^. In brief, the microscope is driven by a mode-locked Ti:Sapphire laser (MaiTai-HP DeepSee, Newport Spectra-Physics, Darmstadt, Germany) and equipped with two fluorescence detection channels and a 20x water immersion objective (W Plan-Apochromat 20x /1.0 DIC M27, Zeiss, Oberkochen, Germany).

**Table 1.** Parts list.

| Part | Description (link) | Company | Item number (RRID, if available) |
| --- | --- | --- | --- |
| ETL | Electrically tunable large aperture lens<br>https://www.optotune.com<br>Specification sheet:<br>https://tinyurl.com/EL-16-40-TC | Optotune, Dietikon, Switzerland | EL-16-40-TC-VIS-5D |
| RL1 | Achromatic Doublet, ARC: 650-1050 nm, F=30mm | Thorlabs | AC254-030-B |
| RL2 | Achromatic Doublet, ARC: 650-1050 nm, F=250mm | Thorlabs | AC254-250-B |
| RL3 | Achromatic Doublet, ARC: 650-1050 nm, F=80mm | Thorlabs | AC254-080-B |
| RL4 | Achromatic Doublet, ARC: 650-1050 nm, F=30mm | Thorlabs | AC127-030-B |
| CM | Cold Light Mirror KS 93 / 45° | Qioptiq Photonics | G380255033 |
| DM1 | Dichroic mirror (custom made) | AHF Analysetechnik AG | F73-063_z400-580-890 |
| BP1 | 510/84 BrightLine HC AOI 0° | AHF Analysetechnik AG | F37-584 |
| BP2 | 610/75 ET Bandpass AOI 0° | AHF Analysetechnik AG | F49-617 |
| ScanM | 2P imaging software running under IGOR Pro | Written by M. Müller (MPI Neurobiology, Martinsried), and T.E. |  |
| IGOR Pro | https://www.wavemetrics.com | Wavemetrics, Lake Oswego, OR | IGOR Pro v6 (SCR_000325) |
| QDSpy | Visual stimulation software<br>https://github.com/eulerlab/QDSpy | Written by T.E, supported by Tom Boissonnet (EMBL, Monterotondo) | - (SCR_016985) |

An ETL with an open aperture of 16 mm (EL-16-40-TC, Optotune, Dietikon, Switzerland) was introduced into the laser path before the scanning unit (Fig. 1a); the optical path after the scanners was left unchanged. Before the ETL, the laser beam is expanded to a diameter of 15 mm using a telescope (T1, a 4f-system with relay lenses RL1,2; *f*_*R*1_=30 mm, *f*_*R*2_=250 mm; for complete parts list, see Table 1). The expanded beam is then reflected perpendicularly to the optical table by a silver mirror towards the horizontally placed ETL, which is housed in a 60 mm cage plate (Thorlabs, Dachau, Germany) using a custom-made adapter (Fig. 1b). After the laser beam passed the ETL, it is narrowed by a second telescope (T2, another 4f-system consisting of relay lenses RL3,4; *f*_*R*3_=80 mm, *f*_*R*4_=30 mm) to a diameter of approx. 5.6 mm, which approx. matches the size of the scanning mirrors. Telescope T2 relays the pupil of the ETL to a conjugate pupil on the scan mirrors. Here, RL4 needs to be positioned precisely at its focal distance and in the centre of the two scan mirrors for correct refocusing of the laser beam. By changing the current supplied to the ETL, it changes its optical power. According to the EL-16-40-TC’s specifications (see link in Table 1), its optical power can be tuned from -2 to +3 dioptres (*f*_*ETL*_ ranging from -500 to 333 mm, for currents of approx. ± 250 mA), rendering the beam divergent or convergent, respectively. Adapting the calculation in Fahrbach et al.^29^, the shift in focal plane (Δ*z*, in [µm]) under the objective lens can be roughly estimated using

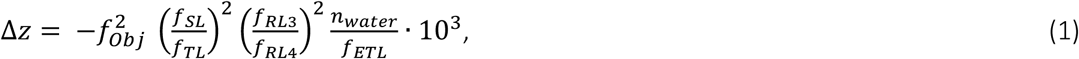

with the focal lengths (in [mm]) of the relay lenses (see above), the scan (*f*_*SL*_=50) and tube lens (*f*_*TL*_=200), and refractive index (*n*_*water*_=1.333). We estimated the objective’s image-side focal length 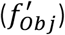 using 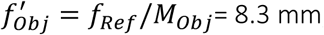, with reference focal length *f*_*Ref*_=165 mm, and magnification *M*_*Obj*_= 20 x. The object-side focal length (*f*_*Obj*_) results from the relationship 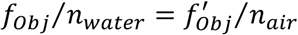, as the objective tip is immersed in solution.

To drive the ETL with our imaging software (ScanM, see below and Table 1), we used custom electronics (designed by R. Berndt, HIH, Tübingen) that translates a voltage signal from one of the analogue-out channels of an PCI 6110 card (NI, Austin, US) controlled by ScanM into a stable current signal. To obtain the relationship between ETL driver input and resulting shift in focal plane, we applied voltage steps of varying amplitude (n=11 amplitudes, n=5 trials), presented in a randomized order, and monitored the shift in focal plane by reading out the z position of the microscope’s motorized scan head. For the used combination of lenses, this resulted in a measured Δ*z* range of +80 and -120 µm (for ETL driving currents of -100 and +100 mA; *cf.* Fig. 2B). Due to technical limitations (i.e. size of the scan mirrors), the Δ*z* range with largely constant laser power spanned approx. 50 µm (*cf.* Fig. 2C), which is sufficient to scan across the entire mouse IPL without adapting the laser power. To characterize the spatial resolution of our system, we measured the point spread function (PSF) of fluorescent beads (170 nm in diameter, *λ*_Emission_=515 nm; P7220; Invitrogen) at different axial planes.

**Figure 2.**
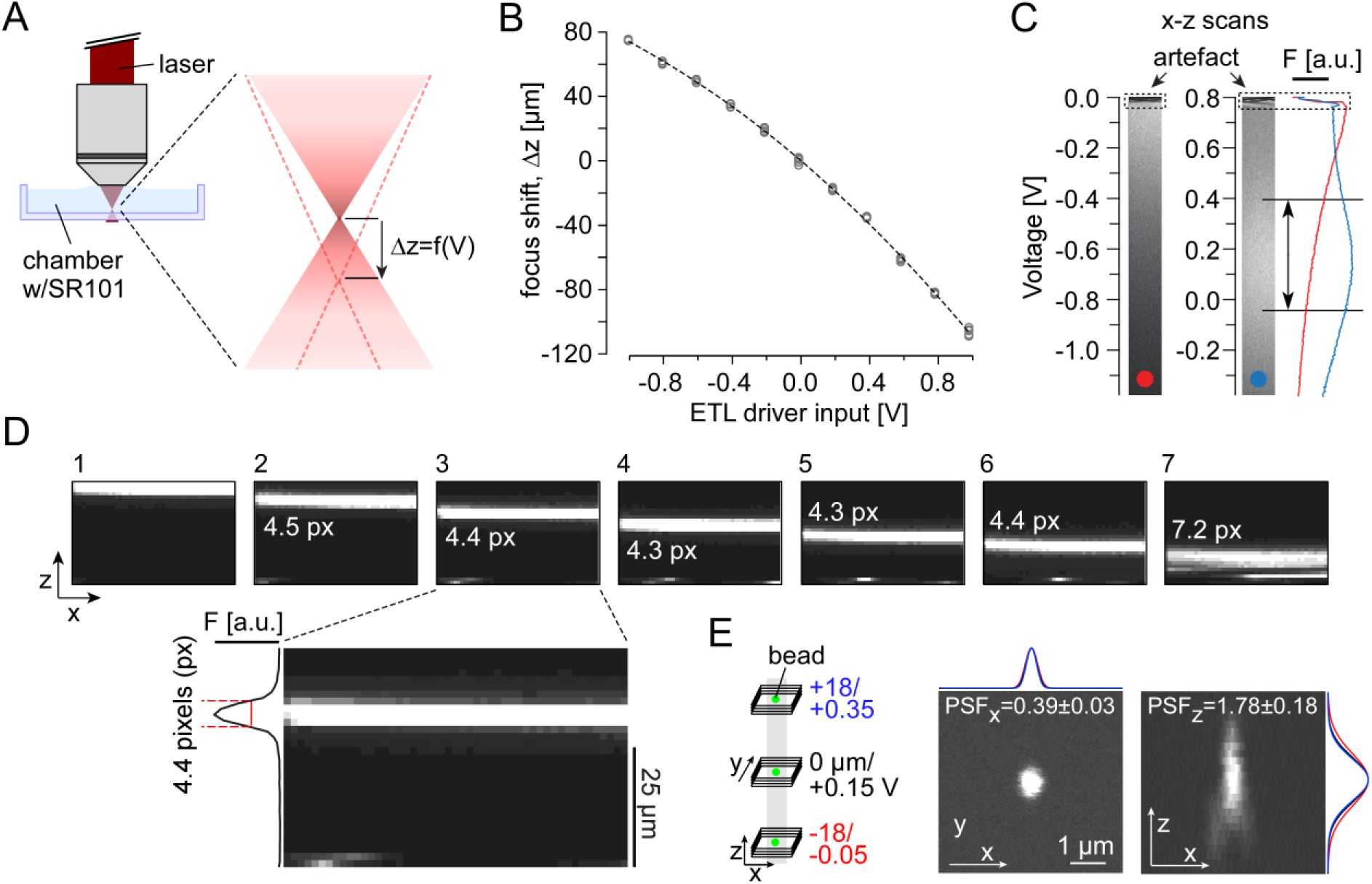
Axial scan properties. A, Illustration of the measurement configuration and the excitation laser’s focus shift (Δ*z*) introduced by the ETL. B, Axial position (measured with the microscope stage motor) as a function of voltage input to ETL driver (circles represent n=5 individual measurements per voltage performed in random sequence; dashed curve represents sigmoidal fit). C, Sulforhodamine 101 (SR101) solution in the chamber was used to measure fluorescence intensity as function of focus shift for two exemplary ETL voltage offsets; x-z scan field (left; 256×256 pixels, 2ms/line; zoom_XY,Z_ = 1.0, 0.8) and mean fluorescence (right). Arrows indicate range (∼40 µm) of near-constant fluorescence. D, Axial x-z scan (64×40 pixels, 2ms/line, zoom_XY,Z_ = 1.0, 1.0, V_Offset_ = 0.15 V) of a 5 μm-thin sheet of fluorescein solution between two coverslips (measured using the microscopes motorized stage) at different z-positions. Inset: Frame 3 with intensity distribution along z-axis; for this scan configuration, the fluorescent band width was 4.4 pixels ± 0.1 (mean ± s.d. for width at half maximum, n=5 measurements), corresponding to a pixel “height” of 1.1 µm. E, Illustration of point spread function (PSF) measurements at three positions (−18 (red), 0 (black), 18 μm (blue)) along the z-axis (right); example images of fluorescent beads (170-nm beads, *λ*_Em, Peak_ = 515 nm; 256×256 pixels, n=60 z-planes, Δz=0.2 µm, zoom_XY_=8) at 0 μm, with mean Gaussian fits (n=3 measurements/plane). PSF_x_ and PSF_z_ indicate the mean ± s.d. across the three axial planes (n=9 measurements; see Table 3).

### Animals and tissue preparation

All animal procedures were approved by the governmental review board (Regierungspräsidium Tübingen, Baden-Württemberg, Konrad-Adenauer-Str. 20, 72072 Tübingen, Germany) and performed according to the laws governing animal experimentation issued by the German Government. For all experiments, we used adult mice of either sex from the following lines: B6;129S6-Chat^tm2(cre)Lowl^J (n=3; ChAT:Cre, JAX 006410), and B6;129P2-*Pvalb*^*tm1(cre)Arbr*^/J (n=6; PV:Cre, JAX 008069). All lines were purchased from The Jackson Laboratory (Bar Harbor, ME). The transgenic lines were crossbred with the Cre-dependent red fluorescence reporter line Gt(ROSA)26Sor^tm9(CAG-tdTomato)Hze^ (Ai9^tdTomato^, JAX 007905) for all experiments. Owing to the exploratory nature of our study, we did not use blinding and did not perform a power analysis to predetermine sample size. For details on the mouse lines and all reagents, see Table 2.

**Table 2.** Mouse lines and reagents.

| Item | Description (link) | Company | Item number |
| --- | --- | --- | --- |
| B6;129S6-Chat<sup>tm2(cre)Lowl</sup>/J | Mice expressing Cre recombinase in cholinergic neurons, without disrupting endogenous Chat expression | Jackson Laboratory | JAX 006410 |
| B6;129P2-Pvalb<sup>tm1(cre)Arbr</sup>/J | Mice expressing Cre recombinase in parvalbumin-expressing, without disrupting endogenous Pvalb expression | Jackson Laboratory | JAX 008069 |
| Gt(ROSA)26Sor<sup>tm9(CAG-tdTomato)Hze</sup> | Mice expressing robust tdTomato fluorescence following Cre-mediated recombination | Jackson Laboratory | JAX 007905 |
| AAV2.hSyn.iGluSnFR.WPRE.SV40 | Viral construct | Penn Vector Core, Philadelphia, PA |  |
| AAV2.7m8.hSyn.iGluSnFR | Viral construct, produced in D. Dalkara's lab with plasmid provided by Jonathan Marvin and Loren Looger, Janelia Research Campus, Ashburn, VA |  |  |
| NaCl | Sodium chloride | VWR | 27810.295 |
| KCl | Potassium chloride | Sigma-Aldrich | P9541 |
| MgCl<sub>2</sub>·6H<sub>2</sub>O | Magnesium chloride hexahydrate | Merck Millipore | 1.05833 |
| NaH<sub>2</sub>PO<sub>4</sub> | Sodium phosphate monobasic | Sigma-Aldrich | S5011 |
| C<sub>6</sub>H<sub>12</sub>O<sub>6</sub> | D-(+)-Glucose | Sigma-Aldrich | G8270 |
| NaHCO<sub>3</sub> | Sodium hydrogen carbonate | Merck Millipore | 1.06329 |
| CaCl<sub>2</sub>·2H<sub>2</sub>O | Calcium chloride dihydrate | Sigma-Aldrich | C3306 |
| C<sub>5</sub>H<sub>10</sub>N<sub>2</sub>O<sub>3</sub> | L-Glutamine | Sigma-Aldrich | G3126 |
| SR101 | sulforhodamine-101 | Invitrogen | S359 |

**Table 3.** Point spread functions (PSF) from n=3 independent measurements at each z position.

| Z position (ETL)<br>[μm] | PSF<sub>x</sub><br>[μm] | PSF<sub>z</sub><br>[μm] |
| --- | --- | --- |
| 18 | 0.43, 0.42, 0.42 | 1.98, 2.02, 2.05 |
| 0 | 0.39, 0.37, 0.38 | 1.78, 1.69, 1.70 |
| -18 | 0.35, 0.33, 0.38 | 1.56, 1.62, 1.67 |

Animals were housed under a standard 12h day-night cycle. For recordings, animals were dark-adapted for >1h, then anaesthetized with isoflurane (Baxter, Deerfield, US) and killed by cervical dislocation. The eyes were removed and hemisected in carboxygenated (95% O_2_, 5% CO_2_) artificial cerebral spinal fluid (ACSF) solution containing (in mM): 125 NaCl, 2.5 KCl, 2 CaCl_2_, 1 MgCl_2_, 1.25 NaH_2_PO_4_, 26 NaHCO_3_, 20 glucose, and 0.5 L-glutamine (pH 7.4). The tissue was then transferred to the recording chamber of the 2P microscope, where it was continuously perfused with carboxygenated (5% CO_2_, 95% O_2_) ACSF at ∼37 °C. The ACSF contained ∼0.1 µM sulforhodamine-101 (SR101, Invitrogen, Carlsbad, US) to reveal blood vessels and any damaged cells in the red fluorescence channel. All procedures were carried out under very dim red (>650 nm) light.

### Intravitreal virus injection

For virus injections, mice were anesthetized with 10% ketamine (Bela-Pharm GmbH & Co. KG, Vechta, Germany) and 2% xylazine (Rompun, Bayer Vital GmbH, Leverkusen, Germany) in 0.9% NaCl (Fresenius, Bad Homburg, Germany). A volume of 1 µl of the viral construct (AAV2.hSyn.iGluSnFR.WPRE.SV40, Penn Vector Core, Philadelphia, PA, or AAV2.7m8.hSyn.iGluSnFR, produced in D. Dalkara’s lab with plasmid provided by Jonathan Marvin and Loren Looger, Janelia Research Campus, Ashburn, VA) was injected into the vitreous humour of both eyes via a Hamilton injection system (syringe: 7634-01, needles: 207434, point style 3, length 51mm, Hamilton Messtechnik GmbH, Höchst, Germany) mounted on a micromanipulator (World Precision Instruments, Sarasota, Germany). Imaging experiments were performed 3 weeks after virus injection.

### Two-photon imaging

We used our microscope’s “green” detection channel (HC 510/84, AHF, Tübingen, Germany) to record iGluSnFR fluorescence, reflecting glutamate signals. In the “red” channel (ET 610/75, AHF), we detected tdTomato to image the ChAT bands (in the ChAT:Cre x Ai9 mice; *cf.* Fig. 3B) or RGC somata (in the PV:Cre x Ai9 mouse), and SR101 fluorescence to measure fluorescence intensities along the z axis (*cf.* Fig. 2C). The laser was tuned to 927 nm for all fluorophores. For data acquisition, we used custom software (ScanM, see Table 1) running under IGOR Pro 6.3 for Windows (Wavemetrics, Lake Oswego, US).

**Figure 3.**
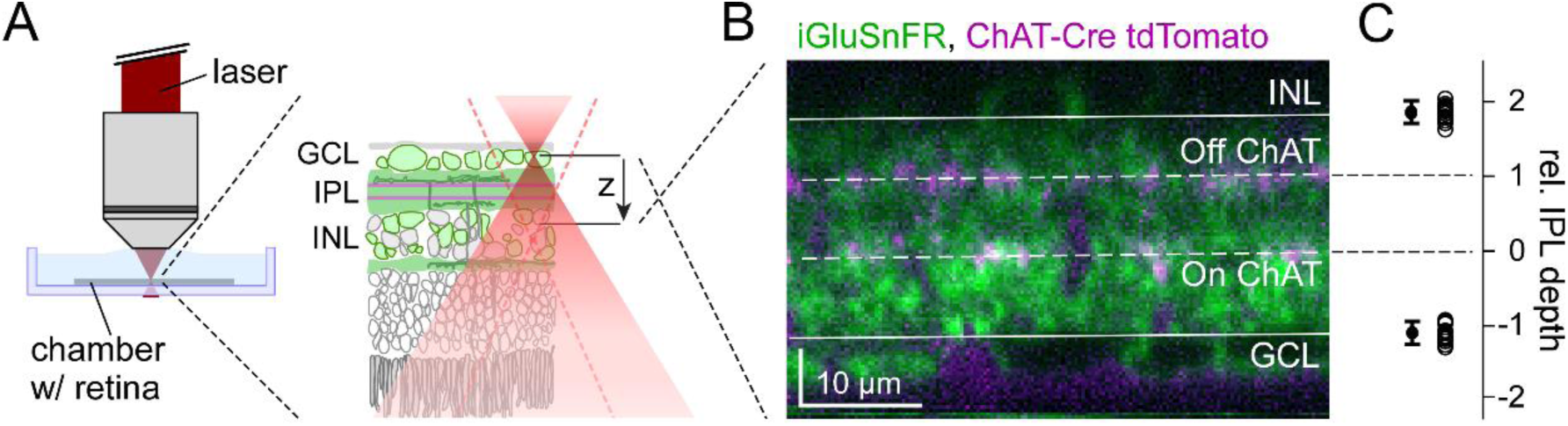
Mapping the inner plexiform layer (IPL). A, Illustration of axial scans in the whole-mount retina of a transgenic mouse expressing tdTomato under the ChAT promotor and iGluSnFR via AAV transduction (Methods). B, Axial x-z scan (256×160 pixels, 2ms/line, zoom_XY,Z_ = 1.5, 0.75) with iGluSnFR expression (green) and ChAT bands (magenta). IPL borders and ChAT bands (solid and dashed lines, respectively) were defined manually (Methods). Note that the retina was flipped from (A), following the convention to show the photoreceptors pointing up. C, IPL border positions relative to ChAT bands (left; INL: 1.9 ± 0.1; GCL: -1.1 ± 0.1; n=3/6/14 mouse/retinas/scans).

### Light stimulation

A modified LightCrafter (DLPLCR4500, Texas instruments; modification by EKB Technology) was focused through the objective lens of the microscope^22,30^. Instead of standard RGB light-emitting diodes (LEDs), it was fitted with a green (576nm) and a UV (390nm) LED for matching the spectral sensitivity of mouse M- and S-opsins^31,32^. To prevent the LEDs from interfering with the fluorescence detection, the light from the projector was band-pass-filtered (ET dual band exciter, 380-407/562-589, AHF) and the LEDs were synchronised with the microscope’s scan retrace^23^. Stimulator intensity was calibrated to range from 0.1×10^3^ (“black” background image) to 20×10^3^ (“white” full field) photoisomerisations P*/s/cone. The light stimulus was centred before every experiment, such that its centre corresponded to the centre of the recording field. Light stimuli were generated using QDSpy, a custom visual stimulation software written in Python 3 (see Table 1). To probe BC function, we presented 3-4 repeats of a “chirp” stimulus in two sizes, local (100 µm in diameter) and global (800 µm); it consisted of a light-On step followed by sinusoidal intensity modulations of increasing frequency and contrasts^7^.

### Data analysis

Data preprocessing was performed in IGOR Pro 6 (Wavemetrics), DataJoint^33^ and Python 3. Regions of Interest (ROIs) were defined automatically by custom correlation-based algorithms in IGOR Pro^7^. First, we estimated a correlation image by correlating the trace of every pixel with the trace of its eight neighbouring pixels and calculating the mean local correlation (ρ_local_). In contrast to previous x-y recordings ^7^, local correlation of neighbouring pixels varied with IPL depth in x-z scans (Fig. 4B) due to differences in iGluSnFR labelling (Fig. 3B) and laser intensity (Fig. 2C). To account for that, an IPL depth-specific correlation threshold (ρ_threshold_) was defined as the 70^th^ percentile of all local correlation values in each z-axis scan line. For every pixel with ρ_local_ > ρ_threshold_ (“seed pixel”), we grouped neighbouring pixels with ρ_local_ > ρ_threshold_ into one ROI. To match ROI sizes with the sizes of BC axon terminals, we restricted ROI diameters (estimated as effective diameter of area-equivalent circle) to range between 1 and 4 μm.

**Figure 4.**
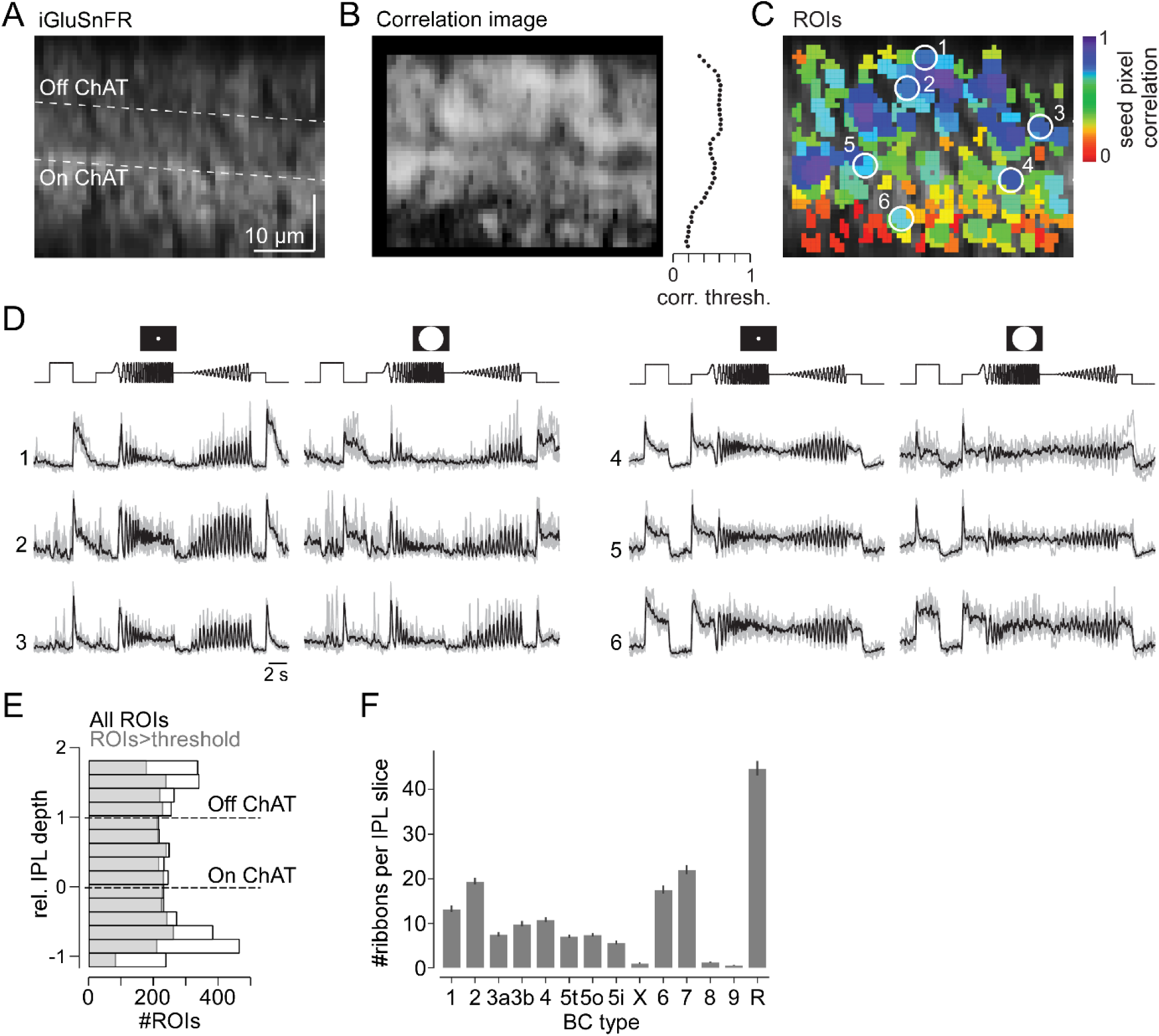
Glutamate imaging in the inner plexiform layer. A, Axial x-z scan (64×56 pixels, 1.6ms/line) of the inner plexiform layer (IPL) in a whole-mount wild-type mouse retina expressing iGluSnFR ubiquitously after AAV-mediated transduction. B, Correlation image (left) and distribution of correlation thresholds across the IPL (right). C, ROIs extracted from scan in (A), pseudo-coloured by seed pixel correlation (for details on ROI extraction, see Methods). D, Glutamate responses to local and global chirp stimulus for exemplary ROIs encircled in (C); Off (left) and On responses (right) are shown. E, Distribution of all ROIs (black, n=5,379) and ROIs that passed our quality threshold (grey, n=3,893; Methods) recorded across the IPL (n=6/8/37 mice/retinas/scans). F, Number of ribbon synapses from different BC types per vertical IPL slice (Methods).

To register each ROI’s depth in the IPL, we determined for each x-z scan the position of the On and the Off ChAT band (Fig. 3; see also Results). In the case of ChAT:Cre mice, ChAT bands could be imaged directly due to their tdTomato fluorescence (Fig. 3B). We found the ChAT band positions can also be reliably determined from the IPL borders, which were determined from the location of iGluSnFR-labelled (or, in PV:Cre x Ai9, tdTomato-labelled) somata in GCL and INL. Here, we first determined the position of the IPL borders relative to tdTomato labelled ChAT bands in a subset of experiments (Fig. 3C). Following conventions^34^, we defined the On and the Off ChAT band position as 0 and 1, respectively. For every ROI, we then estimated the shortest distance to On and Off ChAT bands or IPL borders and expressed ROI IPL depth as a relative value between approx. -1 (GCL border) and +2 (INL border). Next, for every scan field (=batch), ROI depth estimates were corrected using the IPL depth at which the response polarity switched between On and Off BCs. The IPL position of this polarity switch was determined using the first principal component (PC) of the local chirp responses (0.24 ± 0.14, mean ± s.d., n=37 fields) and subtracted from each scan field’s depth estimates. We then added 0.5 to align the IPL centre, i.e. the separation between On and Off BC terminals.

The glutamate traces for each ROI were extracted using the image analysis toolbox SARFIA for IGOR Pro^35^. Then, the traces were synchronised to the light stimuli using time markers that were generated by the stimulation software and acquired during imaging. Finally, we up-sampled the traces to 64 Hz temporal resolution and de-trended them by applying a high-pass Butterworth filter with a cut-off frequency of 0.1 Hz.

### Linear mixed effects model

All modelling was performed using DataJoint and Python 3. Batch effects were first estimated using a series of simple linear mixed effects models, which predicted the expected glutamate release of all ROIs across time (*Y* ∈ ℝ*^NxT^, T* = 64*Hz* · 32*s* = 2048) as a linear function of different predictors that were all one-hot encoded:

1. A model that used only the polarity (*X*_*polarity*_ ∈ ℝ*^Nx2^*) to predict *Ŷ*= *X*_*polarity*_*w*_*polarity*_.Thus, this model computed simply the average of all On and Off ROIs as the weight vector.
2. A model that used only the IPL depth (*X*_*depth*_ ∈ ℝ^*Nx*20^), where the response is estimated non-parametrically with 10 depth bins across the IPL for each polarity (*cf*. Fig 6A), to predict *Ŷ* =*X*_*depth*_*w*_*depth*_.
3. A model that estimated the response of each ROI as a function of the batch (*X*_*batch*_ ∈ ℝ*^Nx2B^*; *B* = *36*, the first batch was left out as reference and to avoid a singular design matrix) where each batch had one predictor per polarity to predict*Ŷ* = *X*_*batch*_*w*_*batch*_.
4. And finally, a combined model (2. and 3.) that predicted the response of each ROI as a function of both batch and depth (*Ŷ*= *X*_*batch*_*w*_*batch*_ *+ X*_*depth*_*w*_*depth*_).

In the last model, we wanted to make sure that any shared variance between the two predictors would be ascribed to the depth predictor – conservatively mistaking nuisance or noise from the batches rather as biological signal than vice versa. To achieve this, we orthogonalized *X*_*batch*_ with respect to *X*_*depth*_, effectively by using *X*_*batch_new*_ = *X*_*batch*_ − *P*_*depth*_*X*_*batch*_, with *P*_*depth*_ denoting the projection matrix onto the subspace defined by *X*_*batch*_.

**Figure 6.**
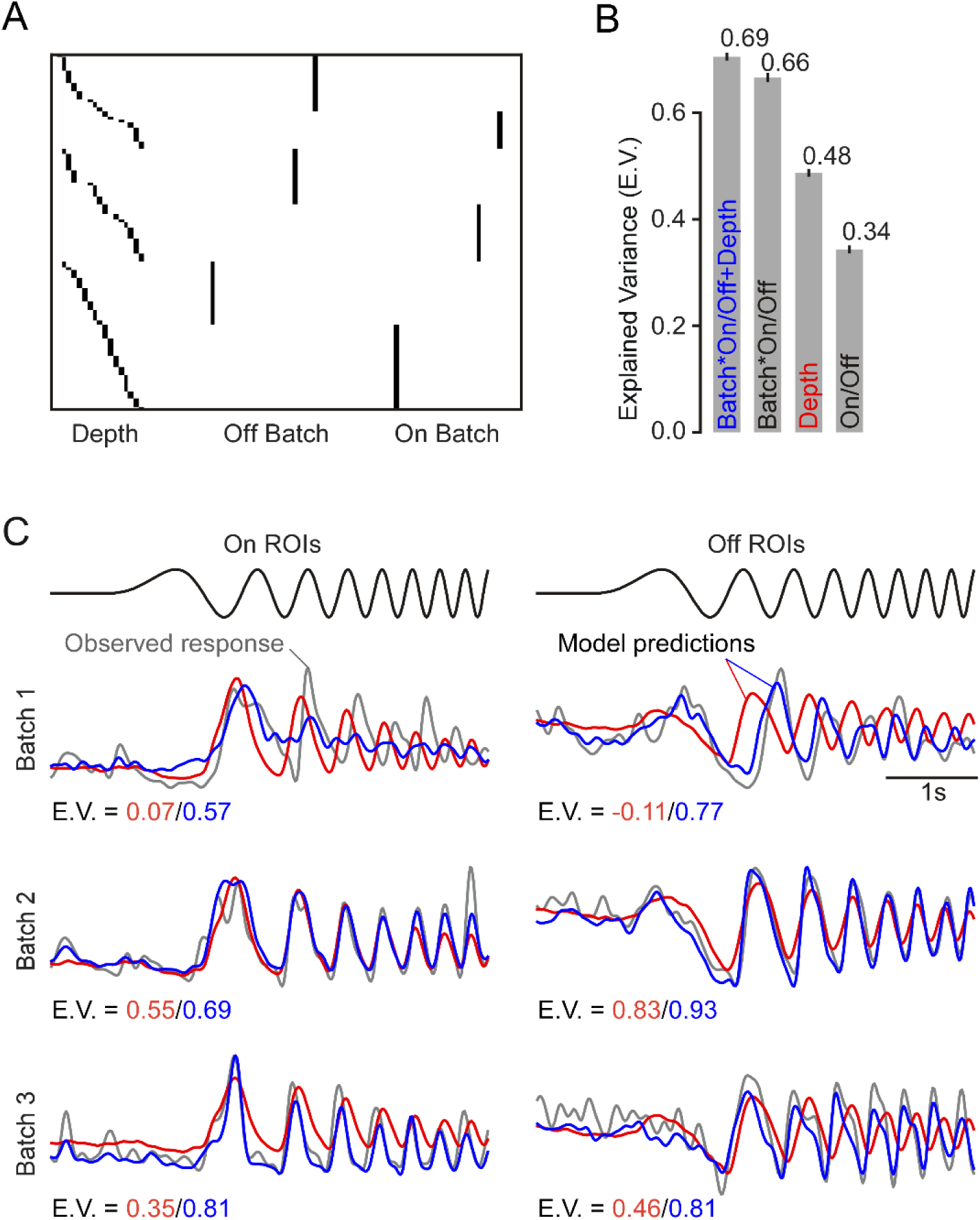
Batch effect estimation using linear models. A, Design matrix with IPL depth and batch specific predictors (example scan fields from Fig. 5). B, Model comparison for On/Off, batch and IPL depth in a linear model fitted to the local chirp response data (same dataset as in Fig. 4A-E). Error bars indicate 2 S.E.M. C, Example traces (grey) for first ROI of each polarity and batch shown in Fig. 5A. Predicted responses from model using IPL depth alone (red) and with an additional batch specific term (blue). E.V., Explained Variance.

### Local chirp encoding model

We also modelled the local chirp responses with different Linear-Nonlinear (LN) encoding models that estimated the finite impulse response, i.e. a temporal receptive field, for each ROI given the chirp input time series. Let *x* denote the chirp stimulus over time, *y*_*i*_ the observed response of ROI *i* ∈ {*1,* … , *N*}. Then we define a temporal filter for this ROI as

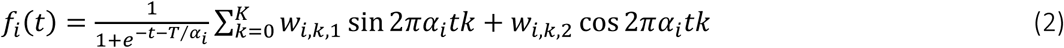

where we set *w*_*i,0,2*_ = 0 to represent the DC component with the sine, *T* = 64 the temporal extent of our filter reaching back 1 s in time (64 Hz), *K* = 21 the highest frequency of the Fourier basis for our kernels (i.e. well below the Nyquist, and reasonably smooth), and *α*_*i*_ a temporal stretching factor. The sigmoidal factor before the sum is a soft-thresholding mask that sets the last part of our filter to 0 to avoid entering the next cycle when *α*_*i*_ > 1. The response is then predicted as

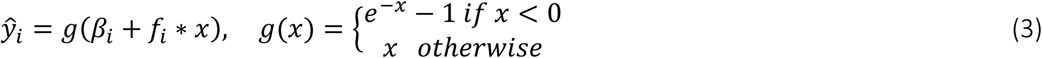

Where *β*_*i*_ is an offset of this ROI and *\** denotes convolution. We fit three models of decreasing flexibility:

1. A model with a separate kernel for each ROI, i.e. fixing *α*_*i*_ = 1 and fitting *w*_*i,k*_ and *β*_*i*_ for all ROIs.
2. A model with one learned shared kernel, i.e. *w*_*i,k*_ = *w*_*j,k*_ ∀*i, j, k*, and one temporal stretch per ROI, i.e. fitting *α*_*i*_ for all ROIs. Additionally, each ROI learned a scalar *a*_*i*_*f*_*i*_ to scale and flip (for On and Off BCs) the shared kernel.
3. Finally, a model with a shared kernel (like model 2.) and a stretch that is a function of the ROI’s depth *d*_*i*_ and its batch *b*_*i*_, i.e. *α*_*i*_ = *h(d_i_, b_i_)*. Firstly, using effectively only the depth and fitting a weight *ξ*_*j*_ for each depth bin *c*_*j*_ (*cf*. Fig 6A) to model the speed of each ROI: 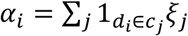. Secondly, the same model but with an additive shift 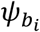 for each batch: 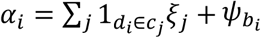. And finally, with an interaction between batch and depth, i.e. separate depth bin weights 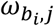 for each batch: 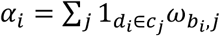.

### Estimating explained variance

For an observed *y* and a predicted *ŷ* response, we estimated the explained variance as

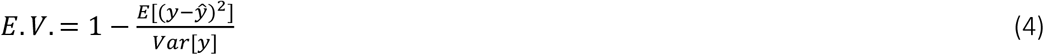

### Estimating BC synapse number in x-z slices

We estimated the average number of synapses (∼ROIs) per BC type to be expected in an x-z scan, which can be considered as a ∼0.5 µm-thick optical section (*cf.* PSF measurements, Table 3). To this end, we used a published EM dataset (e2006, ref^10^) to first determine the volume the axon terminals of each BC type occupies in a 0.5 × 50 x (IPL thickness) µm slice of the IPL. This we did for n=180 non-overlapping slices in the EM stack’s central region, hence limiting the contribution of not fully reconstructed BCs with their soma outside the EM stack. Next, we estimated the number of output synapses (ribbons) per axon terminal volume for each BC type. For this, we calculated the average total axon terminal volume per BC type based on all BCs considered completely reconstructed in the dataset^11^ and divided it by the number of ribbons per type, as reported by Tsukamoto and Omi^36^. Finally, we estimated the number of ribbons (synapses) per slice (x-z scan) and BC type by dividing each BC’s axon terminal volume in a slice by its average axon terminal volume per ribbon (Fig. 4F).

## Results

### Setting up axial scanning

To allow axial scanning of the retina, we inserted an ETL into the optical pathway of the 2P laser of our microscope (Fig. 1). By electrically modulating the optical power of the ETL, the beam of the 2P laser entering the microscope’s objective converges or diverges, resulting in a focus shift along the z-axis of the recording. For the ETL used here, a change in electrical current is transformed into a pressure change, which in turn regulates the lens volume and, thereby, the curvature of the lens surface (Fig. 1A, inset).

When the lens is positioned vertically (w/optical axis parallel to the table), its fluidic core may be slightly deformed by gravitational forces, resulting in a deterioration of its optical properties. The simplest horizontal arrangement would be to place the ETL directly above the objective^25,37^. However, we decided against this possibility for two reasons: First, in this position, the ETL would introduce a focal plane-dependent change in image magnification^25^. Second, if the visual stimulus is coupled into the laser pathway after the scan mirrors – like in our setup (Fig. 1A)^22,30^ – the ETL would also modulate the stimulation plane and spectrally filter the stimulus. The latter is critical if UV stimuli are used: Since prolonged exposure to UV light can degrade the optomechanical properties of the lens, ETLs are typically equipped with a UV-reflecting glass window (for specifications, see Table 1). Therefore, placing the ETL into the stimulus path hampers UV stimulation, but UV stimulation is required to properly drive the mouse retina with its UV-sensitive cone photoreceptors (*λ*_*Peak*_ = 360 nm; as discussed in ref^30^). To avoid these issues, we positioned the ETL horizontally in the pathway that reflects the excitation laser onto the scan mirrors (Fig. 1A, B) (F. Voigt, F. Helmchen, personal communication).

To make use of the full aperture of the ETL, we pre-expanded the laser beam using a telescope system (T1). After the ETL, we used a second telescope system (T2) to refocus the laser beam onto the scanning mirrors. As T2 determines the beam diameter that enters the back aperture of the objective and the z-range of focus shift (together with the objective lens’ magnification), it needs to be defined for the specific microscope and the desired z-range.

First, we evaluated the ETL’s performance using a solution containing the red fluorescent dye SR101 (Fig. 2A). Our combination of custom ETL driver and optical configuration allowed for a practical focus shifting range of approx. 200 µm (Methods). Importantly, the same voltage signal reliably resulted in the same z position of the focal plane (s.d. < 1 µm) and when staying within a limited voltage range (e.g. ±0.4 V), the relationship between voltage input and z position was sufficiently linear (Fig. 2B). However, we consistently observed intensity fluctuations in the first few lines of each x-z scan frame (Fig. 2C,D). This “artefact” can be explained by the fact that for large, rapid changes in voltage input – as it happens, for instance, when jumping back from the last line of a frame to the beginning (retrace) – the ETL’s refractive state does not change instantaneously but requires a few milliseconds to settle^25,29^. We dealt with this problem simply by excluding the first couple of lines (∼10 ms) of each frame from our analysis. In addition, we found that the fluorescence smoothly varied along the z axis (Fig. 2C, left scan). The decrease in fluorescence reflects a loss in laser power, which happens when the ETL changes the beam’s collimation to an extent that the beam becomes too large for the scan mirrors or is partially blocked by down-stream optics. This can be improved by applying an offset voltage to the ETL driver signal, such that the laser intensity peak covers the required z scan range. As the intensity peak is relatively shallow, the IPL of the mouse (∼40 m) can be imaged with almost constant laser intensity.

Next, we tested whether pixel size and spatial resolution remained constant along the z axis of an x-z scan suitable for capturing the complete mouse IPL (e.g. 64×40 pixels corresponding to 50×40 µm). By moving the microscope’s objective lens, we placed a thin fluorescent film (Methods) at different z positions within an x-z scan field (Fig. 2D) and measured the film’s thickness. Apart from the artefact (see above), the recorded thickness of the film remained constant, suggesting that the pixel size is constant along the z axis. Finally, we quantified the spatial resolution of our system by measuring the point spread function (PSF) of fluorescent beads both in the x-y plane and at different z positions (Methods). This was done by first setting one of three z planes (Fig. 2E) using the ETL and then taking image stacks (using the microscope’s motorized stage). The measured PSFs were around 0.4 and 1.8 µm along the x and z axis, respectively, and varied very little for the different ETL planes (Table 3).

In summary, our ETL configuration allows for spatially (nearly) linear, fast axial imaging without detectable loss in spatial resolution.

### Axial scanning in the IPL of the mouse retina

Axial scans were evaluated by imaging light-evoked glutamate release from BC axon terminals (Fig. 3A). After AAV transduction, the glutamate biosensor iGluSnFR^26^ was ubiquitously expressed across the whole retina, including the IPL (Fig. 3B)^7^. As the axon terminals of different BC types stratify at distinct levels of the IPL^10,13,38^, registering IPL depth within the x-z scans is critical. Important landmarks in the IPL are the so-called ChAT bands, which are formed by the dendritic plexi of the cholinergic starburst amacrine cells (SACs)^39^. Accordingly, a commonly used metrics for IPL depth is to define the inner (“On”) band as the origin (=0) and the distance to the outer (“Off”) band as 1 (Fig. 3C) (see also ^7,34^). To relate these positions to IPL borders, we used transgenic mice in which SACs were fluorescently labelled (Fig. 3B; for details, see Methods). We found that the relative distance between ChAT bands and IPL borders was highly consistent across scans and mice (Fig. 3C). Hence, for mice lacking fluorescently labelled ChAT bands, IPL depth can be reliably estimated from the IPL borders.

The ubiquitous expression of iGluSnFR combined with axial scanning allowed sampling of glutamate release at all IPL depths (Fig. 4). To achieve scan rates >10 Hz, we used x-z scans with 64×56 pixels (1.6 ms/line) at a zoom factor that yielded a pixel size of ∼1 µm (Fig. 4A). Regions of interest (ROIs) were based on local image correlations with an IPL depth-dependent threshold (Methods). This ensured ROI placement across the entire IPL (Fig. 4B,C). Subsequently, ROIs were quality-filtered using the reliability of their responses ^7^. Recently, mouse BCs have been functionally characterized by recording BC glutamate release in x-y scans in response to local and global chirp stimuli activating the centre and both centre and surround of the cells’ receptive fields, respectively ^7^. Here, we used the same stimuli to record glutamate traces from BC axon terminals in axial IPL x-z scans (Fig. 4D). As expected from the subdivision of the IPL into On and Off sublamina^19,40^, ROIs located in the outer half displayed increases in glutamate release upon light decrements (Off cells; see ROIs 1–3 in Fig. 4C, D). In contrast, ROIs in the inner part of the IPL showed increased glutamate release upon light increments (On cells; ROIs 4– 6). In addition, response transience varied with IPL depth, with more transient (e.g. ROIs 3 and 4) and sustained BCs (e.g. ROIs 1 and 6) stratifying in the IPL centre and borders, respectively. This observation is in agreement with previous studies demonstrating a spatial segregation of the IPL in transient and sustained BC channels^15,20,21^.

Next, we wanted to evaluate if a single x-z scan – in principle – could capture the full complement of BC types. Overall, ROIs with chirp responses that met our quality criterion^7^ were almost evenly distributed across the IPL (Fig. 4E), suggesting that generally we did not under- or oversample certain IPL levels. However, it has been long known that BC types come at different densities and vary in the size of their axon terminal system^9^. For example, S-cone selective type 9 BCs^11,41^ are low-density and have sparse terminals, whereas rod BCs have small terminals but make up ∼27 % of all BCs in the mouse^9,42^. To estimate the number of ROIs (∼output synapses) the different BC types are expected to contribute to an x-z scan (as in Fig. 4), we calculated the average volume each BC type’s axon terminals occupy in an 0.5 µm (∼PSF_X_) thick slice of the IPL using electron microscopy data^10^ and then determined the mean number of synapses in this volume using published ribbon densities^36^ (Methods; Fig. 4F). This analysis showed that some BC types (i.e. X, 8 and 9) are expected to contribute only very few synaptic terminals to an x-z scan.

Taken together, axial x-z scans allow recording glutamate signals virtually simultaneously across the IPL with similarly high fidelity compared to sequential, hence time-consuming, “traditional” x-y scans. While the functional diversity that can be recovered from individual x-z scans qualitatively resembles that described in an earlier study^7^, reliably capturing signals from BC types with low terminal densities requires integrating data from multiple scans.

### Identification of batch effects

We recorded local chirp light-evoked BC glutamate release from 5,379 ROIs (37 scan fields, 6 mice) across the entire IPL (cf. Fig. 4E). Of those, 3,893 ROIs passed our quality criterion as previously defined in ^7^ and were selected for further analysis. When visually inspecting the data obtained from different recordings, we noticed that the timing of recorded glutamate traces varied systematically across recordings (Fig. 5A). We refer to this variation as “batch effects”, in accordance with similar inter-experimental variability in the RNA sequencing literature (e.g. refs^43–45^).

**Figure 5.**
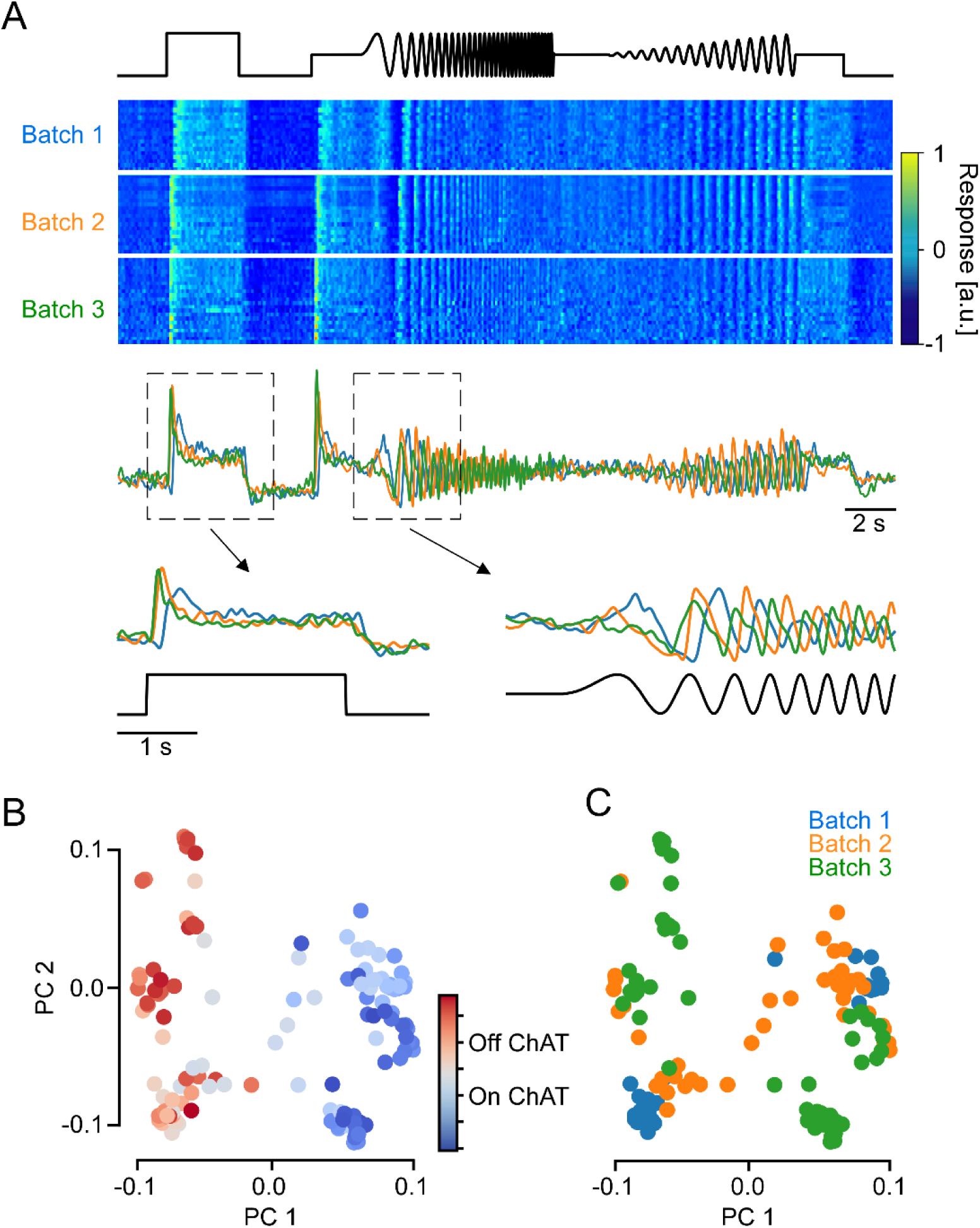
Batch effects in axial x-z scans of the mouse inner plexiform layer (IPL). A, Local chirp responses from ROIs located in the On sublamina of the IPL. From top: time course of chirp stimulus, heat map showing glutamate responses of ROIs from three scan fields (batches), average glutamate responses over ROIs in each batch, and magnified step and frequency responses. B,C, Local chirp responses of ROIs in (A) projected onto their first two principal components (PCs), coloured by IPL depth (B) and batch (C).

In our experiments, the variations between scans may have been caused by experimental factors such as slight temperature fluctuations, as well as differences in light adaptation and/or fluorescence biosensor expression. To investigate the variability in our data, we performed principal component analysis (PCA) on the recorded time-series and inspected the projection onto the first two principal components (PCs; Fig. 5B, C). While On and Off BCs could be distinguished clearly based on the first PC (Fig. 5B), a substantial portion of the variability observed within On and Off BCs seemed to stem from variability across recordings (Fig. 5C). Qualitatively, these batch effects were large enough to be a challenge for recovering the biological differences between cell types within the On and the Off BCs.

We quantified the relative contribution of three factors to the total variance of the observed signal: (1) polarity, i.e. whether the ROI was located in the On or Off sublamina, (2) IPL depth bin in which the ROI was recorded, and (3) the batch (scan field) from which the ROI originated. To this end, we fit a series of linear models (Fig. 6A), each of which included one or more of the three factors (On/Off, IPL depth, batch), and estimated the fraction of variance explained by the models (Fig. 6B,C). The first model, which captured only the polarity, accounted for 34% of the response variance. The second model, which used only IPL depth as a predictor, accounted for 48% of the variance. Note that the first model is a special case of the second one, obtained by splitting the IPL into On and Off sublaminae. As a third model, we used polarity × batch (scan field) ID as a predictor. This model, which amounts to estimating the average trace of On and Off cells in each scan field, accounted for 66% of the variance, substantially outperforming the previous models that only considered the biological source of variation. Finally, adding IPL depth bin as a predictor improved the explained variance only marginally (to 69%).

To summarize, we found that batch effects alone accounted for a larger fraction of the variance than IPL depth (Fig. 6B), which suggests that accounting for such variation can greatly facilitate any analysis of functional differences between BC types beyond On vs. Off.

What is the nature of these batch effects? The most salient difference across the three example batches was a shift in response speed (Fig. 5A). This is especially striking in the response to the chirp’s frequency modulation, where the batch-averaged responses are almost entirely out of phase (Fig. 5A, bottom right). We found the same temporal misalignment in the predictions of our model that considered only IPL depth but ignored the batch effects (Fig. 6C). Comparing the predicted and the recorded traces, we observed that the model was too fast for the first (slow) batch, approximately aligned for the second (medium) batch and too slow for the third (fast) batch. This observation is in line with a previous study that reported systematic differences in the response speed of ganglion cells recorded from different macaque retinae^46^.

A possible explanation for shifts in response speed between experiments may be differences in recording temperature. While we used a closed-loop system to keep the temperature of the tissue at 36°C, small temperature fluctuations in the order of ±1°C cannot be excluded. The temperature coefficient (*Q*_*10*_) of biological reactions, including neural activity, is typically between 2 and 4 (*cf*. ref^47^), therefore, following 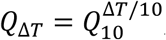, a temperature increase of 1°C may result in a 7 to 15% increase in response speed, which is in the range that we estimated for the batch effects (*cf*. Fig. 8D).

**Figure 8.**
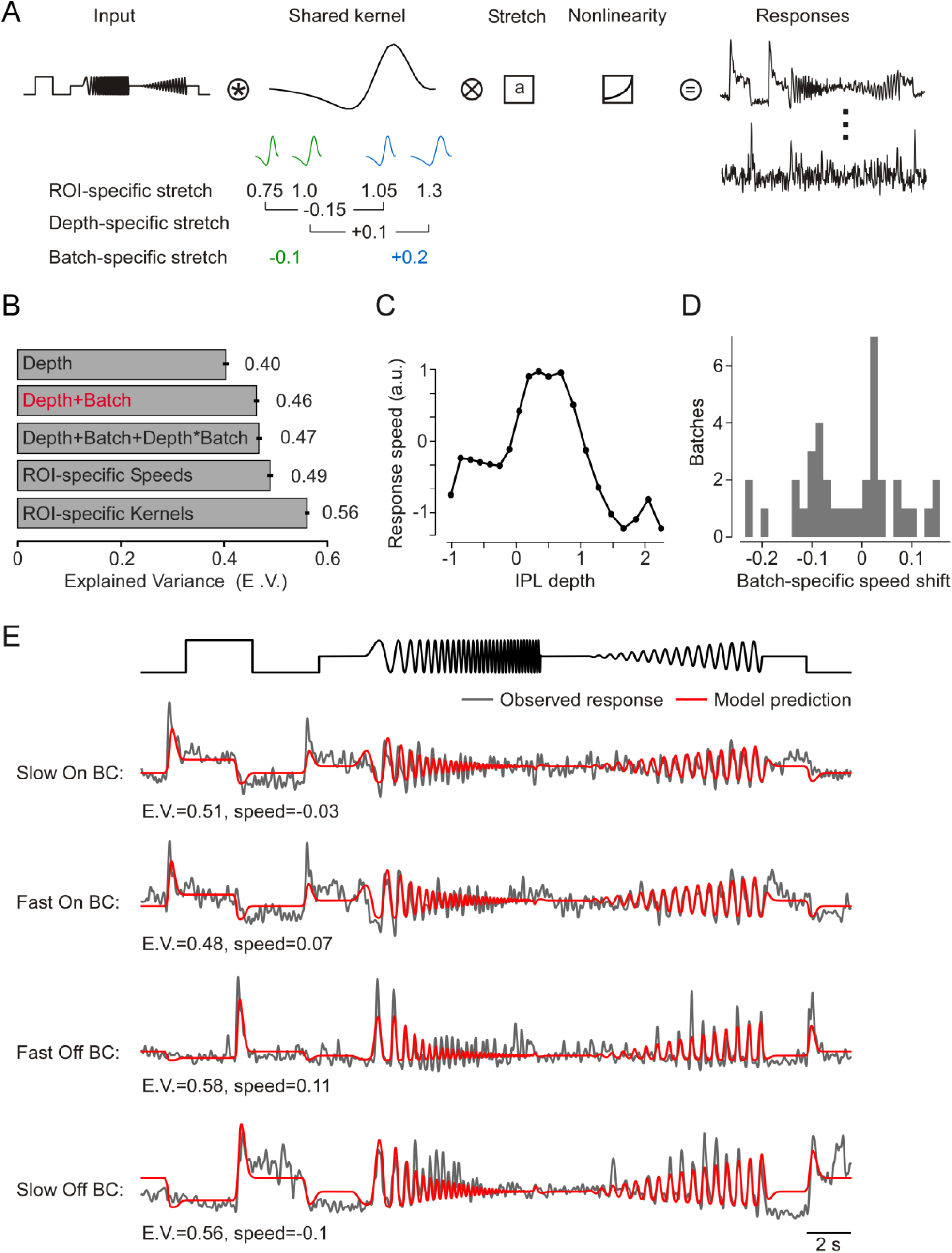
Encoding model comparison. A, Model design: the input time series of the chirp stimulus is convolved with a finite impulse response linear filter which is sign-flipped for On and Off responses and stretched by a factor that is learned for each ROI, then passed through a static nonlinearity (exponential linear unit) and weighted by each ROI to produce the predicted trace. B, Comparison of the different speed models (Methods) learned simultaneously with the encoding model. Error bars indicate 2 S.E.M. C, Speed as a function of IPL depth. D, Distribution over learned batch shifts. E, Observed (black) and predicted (red) traces for four ROIs. E.V., Explained Variance.

### Building batch and IPL depth variations into a shared BC encoding model

To investigate the idea that batch effects effectively result in changes in response kinetics more directly, we fit a linear encoding model and estimated the temporal receptive field kernels of the ROIs in the three example scans shown before. As expected, the temporal kernels showed systematic differences between the three scans that seem to be largely explained by rescaling them in time (Fig. 7A, B). Moreover, within a single batch we could still discern the underlying IPL gradient: ROIs closer to the IPL centre (=lighter colours in Fig. 7C) had their leading edge closer to zero and, hence, responded faster. In addition, central ROIs displayed more biphasic kernels and, hence, responded more transiently.

**Figure 7.**
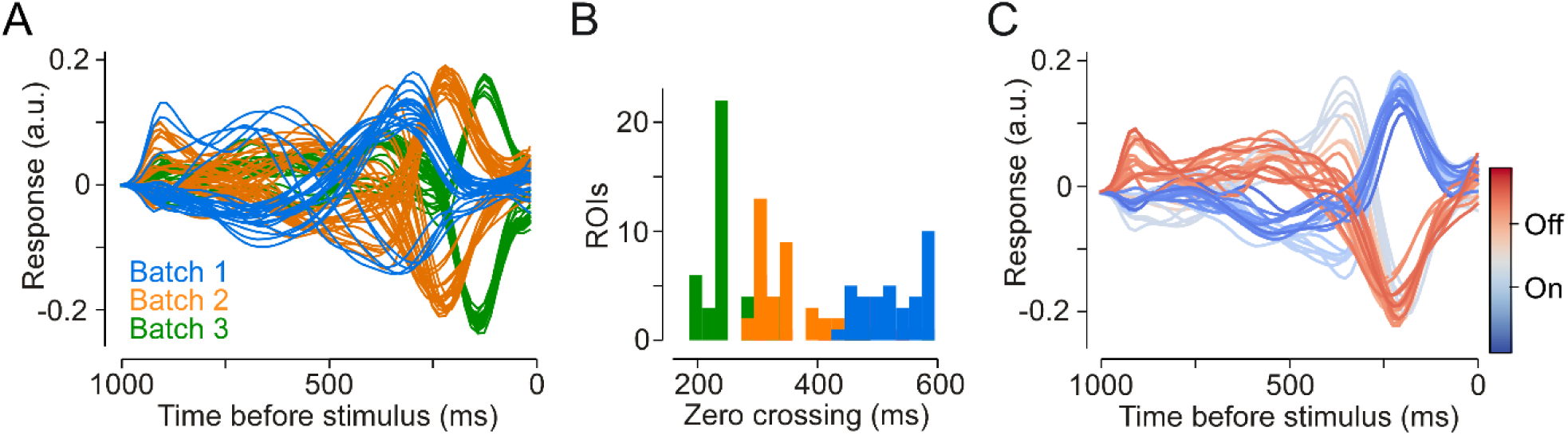
Differences in response speed between batches. A, Learned temporal kernels for all ROIs with E.V. > 0.5. Coloured by batch (same batches as in Fig. 5). B, Zero crossings (after first peak) for all ROIs with E.V. > 0.3. C, All ROIs of the 2^nd^ batch (orange in A, B) with E.V. > 0.3, coloured by IPL depth.

The data presented above suggest that functional differences between individual ROIs can, to a large extent, be accounted for by modelling response speed, and this speed depends on two main factors:(1) laminar location within the IPL and (2) batch effects due to variability between scan fields. We therefore developed a very simple joint encoding model that reduces the functional differences between neurons (here ROIs) to a temporal rescaling of their response kernels (Fig. 8A). The model learned exactly one response kernel that is shared among all neurons and all scan fields. In addition, it allowed for a temporal rescaling of this kernel and a neuron-specific scaling of the response magnitude and polarity-flip (On/Off).

First, we observed that sharing the same kernel shape across all ROIs and only adjusting speed, scale and polarity yielded a predictive performance of 49% explained variance compared to 56% by the model with an individual temporal kernel for each ROI (Fig. 8B). While a 7% difference is not negligible, the difference in complexity between the models is considerable: The latter model used more than 40 parameters for each ROI to specify a response kernel, whereas the former (simplified) model had effectively just 3 parameters to model speed, scale and polarity for each ROI. For the remainder of the paper, we focussed on this simplified model and asked how well we can predict the response speed of individual ROIs based on their IPL depth and a batch-specific speed adjustment.

Next, we tested the effect of additionally constraining this simple model: First, we assumed that the speed of each ROI is only a function of the ROI’s depth in the IPL. This assumption, which meant that all ROIs with the same laminar location share the same response kernel, decreased the predictive performance to 40% explained variance (Fig. 8B). Alternatively, allowing for batch effects by adding a scan field-specific global shift to the speed estimates for all the ROIs in the same scan field, enabled us to capture 46% explained variance. This was similar to the model that allowed each ROI its own speed (49%). Additionally, including an interaction term between IPL depth and batch improved performances only slightly (47%), suggesting that batch variations had similar, approximately additive effects onto the speed across the IPL.

In summary, to the extent that variability between BCs can be modelled by differences in response speed, these speed differences can almost entirely be accounted for (46% vs. 49% explained variance) by laminar location within the IPL and a batch-specific global shift in response speed.

Moreover, with this model we can give a quantitative estimate of the speed gradient across the IPL (Fig. 8C). In line with earlier reports^7,15,20,21^, the BC response speed fell off from the centre of the IPL towards its borders. Notably, between the ChAT bands this speed gradient was nearly symmetrical between On and Off BCs. However, after that the On BCs levelled off and exhibited the same, relatively slow speed until the ganglion cell layer. The Off BCs, by contrast, continued decreasing in their speed almost all the way until the inner nuclear layer.

## Discussion

We implemented fast axial x-z scanning of the whole-mounted mouse retina by equipping a 2P microscope^24^ with an ETL to rapidly shift the focal plane of the laser. We showed that this experimental setup is suitable to record the light-evoked glutamatergic output of BCs almost simultaneously across the complete IPL. Axial scans enabled comparing temporal response properties between IPL strata more directly than “traditional”, time-consuming series of horizontal (x-y) scans. At the same time, x-z scans allowed identifying batch effects, characterized mostly by inter-experimental differences in signal speed. We showed that already batch correction with a simple linear model can improve recovering the characteristic response speed profile across the IPL. Our results indicate that careful consideration of inter-experimental variance is key for extracting functional differences between neurons.

### Techniques for axial scanning

Several technical solutions that enable fast axial scanning in 2P microscopy have been published. Here, two main approaches can be distinguished:

In the first group, a focus change within the sample is realized by moving the objective lens relative to the sample (or vice versa). This has, for instance, been implemented using a piezo to move the objective along the z axis^48^. By coordinating the trajectories of galvo scanners (x-y) and piezo (z), fast volume scanning (i.e. 10 Hz frame rate for a 0.25 mm cube) can be achieved using 3D spiral patterns. While this solution is relatively easy to implement, the inertia of the objective limits the speed with which a focal plane can be selected. In addition, the objective’s movements may introduce vibrations to the sample.

In the second group, focus shifting is achieved by changing the collimation of the laser beam (“remote focussing”) while the objective-to-sample distance remains constant. This strategy eliminates movements close to the sample. A classical solution for remote focussing is to add a reference objective to the laser path to axially displace the focal plane in the sample^49^. Because only a lightweight mirror under this reference objective is moved, this arrangement can reach high axial velocities. However, its superb optical performance is offset by high complexity (i.e. optical alignment; ref^50^) and the costs of a second high-quality objective. In a different approach, an arrangement of inertia-free acousto-optical deflectors (AODs) replaces the galvo scanner and, thus, allows for very fast random-access scanning^25,51–54^. Using a clever AOD arrangement, random-access in 3D is possible^52,55^. Due to the absence of mechanical parts, such a solution enables extremely fast focus changes (<1 ms) at large z ranges (>1 mm). However, AOD-based solutions typically are complex and expensive systems, which usually require substantially more laser power than galvo scanner-based systems.

Comparably fast remote focussing can also be achieved with an ETL, in which the curvature – and hence the focal length – of a liquid lens core is controlled electrically (reviewed in ref^56^). With high-quality inexpensive ETLs becoming available, they offer a cost-efficient and relatively simple way for equipping mechanical scanner-based (or “simple” 2D AOD-based) fluorescent microscopes, including confocal^57^, 2P^25,37,58,59^, and light-sheet systems^29,60^, with fast focussing. In the simplest configuration, the ETL is positioned directly on top of the objective lens^25,37^. However, in this position, shifts in focal plane are accompanied by image scaling in x-y^25^. Also, the ETL’s transmission in the relevant spectral bands – like here, the UV transmission for light stimulation (*cf*. Results) – may need to be considered. By integrating the ETL into the laser path upstream of the scan mirrors^57^, image scaling and (potential) ETL transmission issues are avoided. The additional telescope needed to couple the ETL into the laser path slightly increases complexity but at the same time allows adapting the available z range to the experimental needs.

### *Further improving ETL-based axial scanning in the* ex-vivo *retina*

For x-z scans across the mouse IPL, we used a unidirectional scan mode, where at the end of a frame, the focal position of the excitation laser is shifted back to the first scan line in one ∼50 µm “jump”. This results in the aforementioned artefact in the first few lines of each frame and is caused by fast oscillations in the ETL’s focal power after a rapid change in driving current (see link to specifications, Table 1). The stabilization time of ∼10 ms (for travel distances of ∼50 µm) we observed is consistent with earlier reports^25,61^. For simplicity, we here used scans with more lines, such that the artefact was outside the IPL. Alternatively, optimizing the current trajectory driving the ETL – e.g. by using a steep ramp and “overdriving” the current instead of just a step (*cf*. ref^25^) – may dampen the ETL’s oscillations and, thus, decrease settling time. Also, bidirectional scans that do not require large and rapid changes in z position may reduce such z travel distance-dependent artefacts.

Another potential caveat of an ETL is thermal drift, because driving the lens may slowly heat it up. Since the resistance of the coil that shapes the lens’ core is temperature-dependent, also the current-to-focal power relationship depends on the ETL’s temperature (for details, see ETL specs). We embedded our ETL into a solid adapter ring made from aluminium, which seemed to have kept the ETL’s temperature stable enough, as we did not observe any relevant thermal drifts during the course of a recording. In any case, the ETL model we used features a build-in temperature sensor that can be read out via an I^2^C (Inter-Integrated Circuit) bus connection to monitor the ETL’s temperature.

One consequence of the ETL being positioned upstream of the scan mirrors is a focus dependent change in laser power. Because we needed a relatively small z focus range to scan across the mouse IPL, we applied an offset voltage to the ETL, shifting the shallow peak in laser power to the imaging range. For larger z ranges, one could automatically adjust the laser power with a sufficiently fast modulator (i.e. a Pockels cell) as a function of the ETL’s control signal.

### Identification and removal of batch effects in axial scans of the mouse IPL

For the functional characterization of retinal cell types, our previous studies used data obtained from sequential x-y recordings^7,28^. In the IPL, one disadvantage of this approach is that the sample of BC types in each individual scan greatly varies between scans, and the cells recorded within any one scan will typically share similar response properties. In the middle of the IPL, a third of all BC types can theoretically be recorded in an individual x-y plane (see stratification profiles in refs^10,13,38^), but typical scans will mostly sample 2–3 BC types. In contrast, axial x-z scans established here allow less biased recordings of the glutamate output across the entire IPL, with BC types with very different response properties present in the same scan field. As a result, each scan exhibited a highly stereotypic functional organization of response kinetics across the IPL, as described before^7,15,20,21^. This property greatly facilitated comparison of data obtained by different recordings, which allowed us to detect, quantify and correct for batch-specific variability in BC responses (see below).

In our recent studies characterizing BC and RGC types^7,28^, we did not explicitly correct for batch effects. However, we used other measures to minimize the effect of such inter-experimental variations on clustering. For BC recordings, we estimated a prior probability for cluster allocation for each scan field taken at a specific IPL depth, which was based on the relative axon terminal volume of all BC types at the respective depth (*cf*. Fig. 2c in ref^7^). This is similar to clustering separately within bins of IPL depth, which helps to identify – on average – BC type-specific response signatures. The fact that ROIs from single scan fields were consistently assigned to different functional types suggests that type-specific differences could be resolved despite the variability induced by batch effects. For GCL recordings, we judged response quality based on alpha RGCs^62^, which are easily recognized by their large somata. Only if alpha RGCs displayed their characteristic temporal response profile^62,63^, we included the data. By doing so, we implicitly minimized the variability induced by batch effects. Again, we found that most functional groups were present across experiments, suggesting that experiment-specific speed differences did not induce additional functional clusters. A similar strategy was recently employed in a study on primate RGCs^46^, where the response profile of well-characterized parasol cells was used to normalize and then combine data from different experiments. Thus, if a readily identified and well-calibrated reference exists, batch effects can be reduced by excluding recordings that deviate strongly. However, this approach can only reduce batch effects at the cost of experimental yield. It is therefore not making optimal use of the available data and resolving more subtle differences between cell types will be difficult as they are likely to be masked by batch effects.

Recent evidence indicates that batch effects or inter-retina variability may also comprise subtler effects such as changes in receptive field size, spiking nonlinearity or autocorrelation of cell response^64^. In our data, we observed that temporal effects made large contributions to the batch variability. We therefore focused on modelling them explicitly and found that they recovered most of the predictive performance of much more flexible models.

## Acknowledgments

We thank Aristides Arrenberg, Filip Yaniak, Katharina Rifai and Tom Baden for helpful discussions of the ETL configuration, Gordon Eske for excellent technical assistance, and Jonathan Marvin and Loren Looger for providing viral vectors.

This work was supported by the German Research Foundation (DFG) through Collaborative Research Centre CRC 1233 (project number 276693517) and DFG grants EC 479/1-1, BE5601/2-1, BE5601/4-1, and EU 42/9-1; the European Union’s Horizon 2020 research and innovation programme under the Marie Sklodowska-Curie grant (agreement No 674901); the German Excellence Strategy (EXC 2064 – project number 390727645); the Max Planck Society (M.FE.A.KYBE0004); the German Ministry of Research and Education (01GQ1601/Bernstein Award and 01GQ1002), and the Alexander von Humboldt Foundation (Ref 3.1 - AUS / 1022403 to DP).

The research was also supported by Intelligence Advanced Research Projects Activity (IARPA) via Department of Interior/Interior Business Center (DoI/IBC; contract number D16PC00003). The U.S. Government is authorized to reproduce and distribute reprints for Governmental purposes notwithstanding any copyright annotation thereon. Disclaimer: The views and conclusions contained herein are those of the authors and should not be interpreted as necessarily representing the official policies or endorsements, either expressed or implied, of IARPA, DoI/IBC, or the U.S. Government.

The funders had no role in study design, data collection and analysis, decision to publish, or preparation of the manuscript.

## Author contributions

MB, PB, AE, and TE designed the study; ZZ and AMC designed, implemented and tested the axial recordings with input from DP, KF, and TE; DD produced the iGluSnFR virus; KF performed viral injections with help from TS; KS performed recordings with help from KF; KS performed pre-processing; DK analysed the data with input from PB, AE, KF, TE and LR; CB carried out analysis of EM data; KF, DK, ZZ, and TE prepared the figures; ZZ, DK, KF, AE and TE wrote the manuscript with input from PB and TS.

## References

1. Masland, R. H. The neuronal organization of the retina. Neuron 76, 266–280 (2012).

2. Wässle, H. Parallel processing in the mammalian retina. Nat Rev Neurosci 5, 747–757 (2004).

3. Euler, T., Haverkamp, S., Schubert, T. & Baden, T. Retinal bipolar cells: elementary building blocks of vision. Nat. Rev. Neurosci. 15, 507–519 (2014).

4. Diamond, J. S. Inhibitory Interneurons in the Retina: Types, Circuitry, and Function. Annu. Rev. Vis. Sci. 3, 1–24 (2017).

5. Lukasiewicz, P. D. & Shields, C. R. Different combinations of GABAA and GABAC receptors confer distinct temporal properties to retinal synaptic responses. J. Neurophysiol. 79, 3157–3167 (1998).

6. Eggers, E. D. & Lukasiewicz, P. D. Multiple pathways of inhibition shape bipolar cell responses in the retina. Vis. Neurosci. 28, 95–108 (2011).

7. Franke, K. et al. Inhibition decorrelates visual feature representations in the inner retina. Nature 542, (2017).

8. Ghosh, K., Bujan, S., Haverkamp, S., Feigenspan, A. & Wässle, H. Types of bipolar cells in the mouse retina. J Comp Neurol 469, 70–82 (2004).

9. Wässle, H., Puller, C., Müller, F. & Haverkamp, S. Cone contacts, mosaics, and territories of bipolar cells in the mouse retina. J Neurosci 29, 106–117 (2009).

10. Helmstaedter, M. et al. Connectomic reconstruction of the inner plexiform layer in the mouse retina. Nature 500, 168–174 (2013).

11. Behrens, C., Schubert, T., Haverkamp, S., Euler, T. & Berens, P. Connectivity map of bipolar cells and photoreceptors in the mouse retina. Elife 5, (2016).

12. Shekhar, K. et al. Comprehensive Classification of Retinal Bipolar Neurons by Single-Cell Transcriptomics. Cell 166, 1308-1323.e30 (2016).

13. Kim, J. S. et al. Space-time wiring specificity supports direction selectivity in the retina. Nature 509, 331–6 (2014).

14. Nelson, R., Famiglietti, E. V & Kolb, H. Intracellular staining reveals different levels of stratification for ON- and OFF-center ganglion cells in cat retina. J.Neurophysiol. 41, 472–483 (1978).

15. Baden, T., Berens, P., Bethge, M. & Euler, T. Spikes in mammalian bipolar cells support temporal layering of the inner retina. Curr. Biol. 23, 48–52 (2013).

16. DeVries, S. H. Bipolar cells use kainate and AMPA receptors to filter visual information into separate channels. Neuron 28, 847–856 (2000).

17. Li, W. & DeVries, S. H. Bipolar cell pathways for color and luminance vision in a dichromatic mammalian retina. Nat Neurosci 9, 669–675 (2006).

18. Breuninger, T., Puller, C., Haverkamp, S. & Euler, T. Chromatic bipolar cell pathways in the mouse retina. J Neurosci 31, 6504–6517 (2011).

19. Euler, T., Schneider, H. & Wässle, H. Glutamate responses of bipolar cells in a slice preparation of the rat retina. J.Neurosci. 16, 2934–2944 (1996).

20. Borghuis, B. G., Marvin, J. S., Looger, L. L. & Demb, J. B. Two-photon imaging of nonlinear glutamate release dynamics at bipolar cell synapses in the mouse retina. J Neurosci 33, 10972–10985 (2013).

21. Roska, B. & Werblin, F. Vertical interactions across ten parallel, stacked representations in the mammalian retina. Nature 410, 583–587 (2001).

22. Euler, T. et al. Eyecup scope—optical recordings of light stimulus-evoked fluorescence signals in the retina. Pflügers Arch. - Eur. J. Physiol. 457, 1393–1414 (2009).

23. Euler, T., Franke, K. & Baden, T. Studying a Light Sensor with Light: Multiphoton Imaging in the Retina. Prepr. 2019030244 (2019).

24. Denk, W., Strickler, J. H. & Webb, W. W. Two-Photon Laser Scanning Fluorescence Microscopy. Science (80-.). 248, 73–76 (1990).

25. Grewe, B. F., Voigt, F. F., van’t Hoff, M. & Helmchen, F. Fast two-layer two-photon imaging of neuronal cell populations using an electrically tunable lens. Biomed. Opt. Express 2, 2035–46 (2011).

26. Marvin, J. S. et al. An optimized fluorescent probe for visualizing glutamate neurotransmission. Nat Methods 10, 162–170 (2013).

27. Werblin, F., Roska, B. & Balya, D. Parallel processing in the mammalian retina: lateral and vertical interactions across stacked representations. Prog Brain Res 131, 229–238 (2001).

28. Baden, T. et al. The functional diversity of retinal ganglion cells in the mouse. Nature 529, 345–350 (2016).

29. Fahrbach, F. O., Voigt, F. F., Schmid, B., Helmchen, F. & Huisken, J. Rapid 3D light-sheet microscopy with a tunable lens. Opt. Express 21, 21010 (2013).

30. Franke, K. et al. An arbitrary-spectrum spatial visual stimulator for vision research. bioRxiv 649566 (2019). doi:10.1101/649566

31. Jacobs, G. H., Neitz, J. & Deegan, J. F.. 2. d. Retinal receptors in rodents maximally sensitive to ultraviolet light. Nature 353, 655–656 (1991).

32. Baden, T. et al. A tale of two retinal domains: Near-Optimal sampling of achromatic contrasts in natural scenes through asymmetric photoreceptor distribution. Neuron 80, 1206–1217 (2013).

33. Yatsenko, D. et al. DataJoint: managing big scientific data using MATLAB or Python. bioRxiv (Cold Spring Harbor Labs Journals, 2015). doi:10.1101/031658

34. Sümbül, U. et al. A genetic and computational approach to structurally classify neuronal types. Nat. Commun. 5, 3512 (2014).

35. Dorostkar, M. M., Dreosti, E., Odermatt, B. & Lagnado, L. Computational processing of optical measurements of neuronal and synaptic activity in networks. J Neurosci Methods 188, 141–150 (2010).

36. Tsukamoto, Y. & Omi, N. Classification of Mouse Retinal Bipolar Cells: Type-Specific Connectivity with Special Reference to Rod-Driven AII Amacrine Pathways. Front. Neuroanat. 11, 92 (2017).

37. Annibale, P., Dvornikov, A. & Gratton, E. Electrically tunable lens speeds up 3D orbital tracking. Biomed. Opt. Express 6, 2181 (2015).

38. Greene, M. J., Kim, J. S. & Seung, H. S. Analogous Convergence of Sustained and Transient Inputs in Parallel On and Off Pathways for Retinal Motion Computation. Cell Rep. 14, 1892–1900 (2016).

39. Vaney, D. I. ‘Coronate’ amacrine cells in the rabbit retina have the ‘starburst’ dendritic morphology. Proc R Soc L. B Biol Sci 220, 501–508 (1984).

40. Werblin, F. S. & Dowling, J. E. Organization of the retina of the mudpuppy, Necturus maculosus. II. Intracellular recording. J Neurophysiol 32, 339–355 (1969).

41. Haverkamp, S. et al. The primordial, blue-cone color system of the mouse retina. J Neurosci 25, 5438–5445 (2005).

42. Strettoi, E. & Volpini, M. Retinal organization in the bcl-2-overexpressing transgenic mouse. J. Comp. Neurol. 446, 1–10 (2002).

43. Ritchie, M. E. et al. limma powers differential expression analyses for RNA-sequencing and microarray studies. Nucleic Acids Res. 43, e47–e47 (2015).

44. Haghverdi, L., Lun, A. T. L., Morgan, M. D. & Marioni, J. C. Batch effects in single-cell RNA-sequencing data are corrected by matching mutual nearest neighbors. Nat. Biotechnol. 36, 421–427 (2018).

45. Johnson, W. E., Li, C. & Rabinovic, A. Adjusting batch effects in microarray expression data using empirical Bayes methods. Biostatistics 8, 118–127 (2007).

46. Rhoades, C. E. et al. Unusual Physiological Properties of Smooth Monostratified Ganglion Cell Types in Primate Retina. Neuron 103, 658-672.e6 (2019).

47. Hille, B. Ion channels of excitable membranes. (Sinauer Associates, Inc, 2001).

48. Göbel, W., Kampa, B. M. & Helmchen, F. Imaging cellular network dynamics in three dimensions using fast 3D laser scanning. Nat. Methods 4, 73–9 (2007).

49. Botcherby, E. J., Juskaitis, R., Booth, M. J. & Wilson, T. Aberration-free optical refocusing in high numerical aperture microscopy. Opt. Lett. 32, 2007–9 (2007).

50. Corbett, A. D. et al. Quantifying distortions in two-photon remote focussing microscope images using a volumetric calibration specimen. Front. Physiol. 5, 384 (2014).

51. Katona, G. et al. Fast two-photon in vivo imaging with three-dimensional random-access scanning in large tissue volumes. Nat. Methods 9, 201–8 (2012).

52. Duemani Reddy, G., Kelleher, K., Fink, R. & Saggau, P. Three-dimensional random access multiphoton microscopy for functional imaging of neuronal activity. Nat. Neurosci. 11, 713–720 (2008).

53. Vučinic, D. & Sejnowski, T. J. A Compact Multiphoton 3D Imaging System for Recording Fast Neuronal Activity. PLoS One 2, e699 (2007).

54. Fernández-Alfonso, T. et al. Monitoring synaptic and neuronal activity in 3D with synthetic and genetic indicators using a compact acousto-optic lens two-photon microscope. J. Neurosci. Methods 222, 69–81 (2014).

55. Cotton, R. J., Froudarakis, E., Storer, P., Saggau, P. & Tolias, A. S. Three-dimensional mapping of microcircuit correlation structure. Front. Neural Circuits 7, 151 (2013).

56. Blum, M., Büeler, M., Grätzel, C. & Aschwanden, M. Compact optical design solutions using focus tunable lenses. in International Society for Optics and Photonics 81670W (2011). doi:10.1117/12.897608

57. Jabbour, J. M. et al. Optical axial scanning in confocal microscopy using an electrically tunable lens. Biomed. Opt. Express 5, 645 (2014).

58. Chen, J. L., Pfäffli, O. A., Voigt, F. F., Margolis, D. J. & Helmchen, F. Online correction of licking-induced brain motion during two-photon imaging with a tunable lens. J. Physiol. 591, 4689–98 (2013).

59. Bar-Noam, A. S., Farah, N. & Shoham, S. Correction-free remotely scanned two-photon in vivo mouse retinal imaging. Light Sci. Appl. 5, e16007–e16007 (2016).

60. Mickoleit, M. et al. High-resolution reconstruction of the beating zebrafish heart. Nat. Methods 11, 919–22 (2014).

61. Nakai, Y. et al. High-speed microscopy with an electrically tunable lens to image the dynamics of in vivo molecular complexes. Rev. Sci. Instrum. 86, 013707 (2015).

62. Krieger, B., Qiao, M., Rousso, D. L., Sanes, J. R. & Meister, M. Four alpha ganglion cell types in mouse retina: Function, structure, and molecular signatures. PLoS One 12, e0180091 (2017).

63. van Wyk, M., Wässle, H. & Taylor, W. R. Receptive field properties of ON- and OFF-ganglion cells in the mouse retina. Vis. Neurosci. 26, 297–308 (2009).

64. Shah, N. et al. Learning variability in the neural code of the retina. in Cosyne Abstracts (2019).

